# Curcumin combined with Glu-GNPs enhancing radiosensitivity of transplanted tumor of MDA-MB-231-luc cells in nude mice

**DOI:** 10.1101/2020.07.07.191221

**Authors:** Mengjie Li, Tingting Guo, Yujian Wu, Jiayi Lin, Ke Yang, Chenxia Hu

## Abstract

Breast cancer is the first cause of cancer death in women all over the world. And the morbidity of breast cancer has been increasing in recent years. Nowadays, the treatment of breast cancer is still a problem. Radiotherapy resistance is one of the important factors leading to poor prognosis of clinical treatment. Curcumin, a plant polyphenol from Curcuma longa, has antitumor activity and inhibition of tumor recurrence and metastasis. However, the influence of curcumin on radiosensitivity of breast carcinoma xenografts, the synergistic effect and possible mechanism of curcumin and Glucose-Gold nanoparticles (Glu-GNPs) in nude mice in vivo need to be further explored. The model of transplanted tumor in nude mice was established and sensitized by intravenous injection of curcumin and Glu-GNPs. The results showed that curcumin can significantly inhibit the growth of tumor, and has no obvious effect on the body weight of nude mice, and Curcumin, Glu-GNPs and X-ray irradiation treatment alone or in combination could reduce the expression of *VEGF* and *HSP90* mRNA and protein compared with model group in tumor tissue. The inhibition of irradiation resistance may be related to inhibiting the synthesis of *VEGF* and *HSP90*. This study suggested that curcumin and Glu-GNPs have the radiosensitization effect in vivo, which provides an experimental basis for the clinical development of green radiosensitizers.

## Introduction

According to the data published by the International Cancer Research Institute in 2018, breast cancer has become the second leading cause of cancer related death and the first one for women all over the world ^[1]^. It is expected that the morbidity and mortality will still increase significantly in the next few years ^[2]^. Nowadays, the treatment of breast cancer mainly includes surgery, chemotherapy and radiotherapy ^[3-4]^. Although great progress has been made in the early breast cancer treatment, there is no effective strategies to deal with breast cancer metastasis. Establishing a nude mouse model of breast cancer xenograft and studying the important related genes in breast cancer growth and development, is an important breast cancer research method, and plays a key role in studying the etiology, pathogenesis, effective prevention, diagnosis and treatment, drug screening and rehabilitation of breast cancer ^[5]^. In recent years, bioluminescence imaging system has become a sensitive research method in animal model. It can monitor the growth and metastasis of tumor in real time. In addition, it can be accurately detected in vivo without killing mice ^[6]^. Zhang et al ^[7]^ found that the results of in vivo imaging and pathological detection are consistent, but compared with HE staining, the growth and metastasis of breast cancer can be more directly and clearly observed using in vivo imaging system. In vivo imaging method is applied to study transplanted tumor in mice. Compared with traditional measurement method, in vivo imaging shows scientific methods to measure tumor growth more accurately, which greatly reduces the error of artificial measurement.

The development of Gold nanoparticles (GNPs) can be traced back to the 16th century. GNPs, a type of nanomaterial approved for clinical trials by FDA in the US, had become a hot area in scientific research ^[8]^. It was reported that when GNPs received X-ray irradiation, the ionizing ability of the radiation to tumor cells was enhanced, which promoted tumor cell damage and apoptosis ^[9-12]^. Our previous study suggested that Glu-GNPs assimilated in MCF-7 adherent cells and THP-1 suspension cells were easier than GNPs and enhanced the cancer killing of THP-1 cells 20% more than GNPs ^[13]^. It pointed that glucose tagging may be a correct way to promoting the uptake of GNPs in tumor cells, as tumor cells can take up more glucose than normal cells. In addition, it was previously demonstrated that these nanometal particles enhanced killing effects on tumor cells without increasing damage to the surrounding normal tissues in a mouse model, thereby reducing the adverse effects of radiotherapy ^[14-15]^.

In recent years, scientific research has found that traditional Chinese materia medica has unique advantages and efficacy in treating tumors. Curcumin, a plant polyphenol extracted from the rhizome of *Curcuma longa*, is the main ingredient of curry, with the characteristics of high efficiency, safety and low toxicity ^[16]^. It was reported that curcumin has killing and therapeutic effects on many kinds of cancer including lung cancer ^[17]^, gastric cancer ^[18]^ and breast cancer ^[19]^ in vitro and in vivo. Hu Shan, et al ^[20]^ found that curcumin can inhibit the proliferation of various breast cancer cells such as MDA-MB-231, MCF-7, MDA-MB-468 and T47D, which also induced cell apoptosis. Palange’s study showed that curcumin treatment can limit the metastatic potential of breast cancer cell lines, possibly by altering the expression of related adhesion molecules ^[21]^. These findings suggested that curcumin may have important research significance in the treatment of breast cancer. However, it reminds to be studied whether curcumin enhances radiosensitivity effect and curcumin has synergistic effect with Glu-GNPs. And the mechanism of curcumin on breast cancer is not clear.

In this study, we used the cell line MDA-MB-231-luc to establish subcutaneous transplant tumor model. The tumor-bearing mice were treated with curcumin, Glu-GNPs and irradiation alone or in combination for 3 weeks. The tumor volume and body weight of tumor-bearing mice were measured regularly. Tumor growth was monitored by in-vivo imaging system. Our results showed that curcumin may inhibit tumor growth in nude mice and reduce tumor fluorescence intensity but has no obvious effect to the body weight. Further study suggested that curcumin and Glu-GNPs along with X-ray irradiation can down-regulated the mRNA and protein expression of *VEGF* and *HSP90*. These data showed that curcumin may be a potential drug for breast cancer treatment.

## Materials and methods

### Reagents and antibodies

Curcumin(Sigma-Aldrih, Merck KGaA Darmstadt, Germany; Purity≥80%)was dissolved in corn oil with 2% dimethyl sulfoxide(DMSO).The concentration of Glu-GNPs(Beijing Dk Nanotechnology Co, Ltd, Beijing, China) is 1 mg/mL. The primary antibodies [*β-actin* (#4970), *HSP90* (#4877)] were obtained from Cell Signaling Technology, *VEGF* (ab52917) from Abcam and the secondary antibodies goat anti-rabbit and goat anti-mouse were obtained from Boster (Boster Bioengineering Co, Ltd, Wuhan).

### Cell culture

Human breast cancer cell line MDA-MB-231 cells labeled with firefly luciferase (MDA-MB-231-luc) was obtained from Shanghai Soft Top Biological Company and cultured in Modified Eagle’s Medium (MEM) supplemented with 10% Fetal Bovine Serum(FBS),100U/ml penicillin and 100μg/ml streptomycin(all from Gibco, USA) at 37°Cin a humidified incubator with an atmosphere of 5% CO_2_ (Thermo fisher scientific). The cells were passaged at a density of 2.0×10^5^ cells in a 25cm^2^ flask or cryopreserved at a density of 2.0×10^6^/mL.

### Detection of MDA-MB-231-luc fluorescence intensity

The fluorescence intensity of MDA-MB-231-luc was measured by in-vivo imaging system. Different number of cells(1.25×10^4^, 2.5×10^4^, 5×10^4^ and 1×10^5^ respectively)were inoculated into the black opaque 96-well plate for 4 hours to make the cells attach to the well. According to the operation of the Firefly-Luciferase assay kit, the cells were lysed with lysate and then added with luciferase substrate. Their luminescence was detected in-vivo imaging system immediately.

### Establishment of subcutaneous transplanted tumors in nude mice

Female *BALB/c-nu/nu* mice aged 4-6 weeks were purchased from Guangzhou University of Chinese Medicine and the scientific research was passed the animal ethics examination. The nude mice were raised adaptively for 7 days and after that they were randomly divided into groups including control group, model group(Model), cisplatin group(Cis), curcumin group(Cur), irradiation group(IR), curcumin combined with Glu-GNPs group (Cur+Glu-GNPs) and curcumin combined Glu-GNPs along with irradiation group (Cur+Glu-GNPs+IR) (5-8 mice per group). All nude mice except the control group were inoculated 0.1mL cell suspension containing 1.5×10^7^ MDA-MB-231-luc cells in phosphate-buffer saline (PBS) and Matrigel (1:1) into the second pair of subcutaneous mammary glands on the left. Nude mice in control group were inoculated with 0.1mL suspension of PBS and Matrigel (1:1). After 3-7 days, tumors formed in the inoculated site of nude mice except the control group. When the diameter of transplanted tumor was greater than 5 mm, the model was successfully established. The tumor volume was measured every 3 days and the body weight of mice was done every week.

### Tumor-bearing mice treated with drugs and irradiation

The calculation method of tumor volume is as follows: V=ab^2^/2(a, length; b, short diameter).When the volume of the tumor reached 100-200 mm^3^, the tumor-bearing mice of Cur group were treated with curcumin (100 mg/kg curcumin dissolved in corn oil contained 2% DMSO), Model group (corn oil contained 2% DMSO), IR group (corn oil contained 2% DMSO), and Cur+IR group (100 mg/kg curcumin dissolved in corn oil contained 2% DMSO) by intraperitoneal injection every other day, and Glu-GNPs+IR group (4 mg/kg Glu-GNPs suspension) by caudal vein injection weekly, and Cisplatin group (3 mg/kg cisplatin dissolved in normal saline)by intraperitoneal injection every three days, or Cur+Glu-GNPs+IR (100 mg/kg curcumin by intraperitoneal injection every other day and 4 mg/kg Glu-GNPs by caudal vein injection weekly). The treatment lasted for 3 weeks. After that, the tumor-bearing mice in IR, Glu-GNPs+IR and Cur+Glu-GNPs+IR group were treated with X-ray irradiation (10 Gy) by X-Ray Irradiation Cabinet (MultiRad 225, Faxitron Bioptics, LLC. America). All tumor tissue was extracted from tumor-bearing mice and weighted on the 24th of treatment. One part of the tumor tissue was fixed by soaking in 4% polyformaldehyde solution at 4°C for HE staining and immunohistochemistry assay. The other part of the tumor was transferred into liquid nitrogen and stored in the refrigerator at −80°C for qRT-PCR detection.

### Tumor growth was monitored using in-vivo imaging system

After treatment for 24 days, the tumor-bearing mice were injected 150mg/kg luciferase substrate (sciencelight, shanghai, China) intraperitoneally in advance. Mice were anesthetized by inhaling isoflurane during observation. The mice were lying on their sides, and the tumor was exposed. The fluorescence intensity in all groups of mice was detected and recorded 10 minutes after the injection by using in-vivo bioluminescence imaging system.

### HE staining

Tumor tissue samples were fixed in 4% paraformaldehyde at 4°C for 48 hours. After that, the samples were dehydrated, embedded, paraffin sectioned and stained according to the general process. The malignant degree of tumor cells in each group was observed under microscope after sealed with neutral gum.

### Immunohistochemical analysis

*VEGF* (*VEGF* Rabbit mAb, 1:50) and *HSP90* (*HSP90* Rabbit mAb, 1:50) expression were detected by immunohistochemistry. Tumor tissue samples fixed in 4% polyformaldehyde solution were sliced with the paraffin-sectioning machine, stained with an immunohistochemistry kit (Bioss, China), and then sealed with neutral gum after dehydration of gradient ethanol solution and transparent treatment of xylene. Positive cells were randomly selected five areas and counted under 400× optical microscopy.

### Quantitative Real-Time Polumeease Chain Reaction(qRT-PCR)

Total RNA was extracted using trizol (Takara, Japanese) and cDNA synthesis using Takara RNA Purification Kit according to the manufacturer’s instructions. The amplification procedure was as follows: the first step is 95 ° C for 3 minutes, the second step is at 95 ° C for 5 seconds, and 60 ° C for 30 seconds for 40 cycles, and the last step ends with the Melt Curve procedure. The relative expression was evaluated following the relative quantification equation, 2^-ΔΔCT^.qPCR was performed by using the following primers:

*ACTB*:forward:5’*GTGGCCGAGGATTTGATTG*3’;

reverse:5’*CCTGTAACAACGCATCTCATATT*3’

*GAPDH*:forward:5’*AGCCACATCGCTCAGACAC*3’;

reverse:5’*GCCCAATACGACCAAATCC*3’

*VEGF*:forward:5’*TAGAGTACATCTTCAAGCCGTC*3’;

reverse:5’*CTTTCTTTGGTCTGCATTCACA*3’

*HSP90*:forward:5’*ACGAAGCATAACGACGATGAGCAG*3’;

reverse:5’*CCATTGGTTCACCTGTGTCAGTCC*3’

### Statistical Analysis

The data of multiple groups were analyzed by one-way ANOVA, and subsequent analysis was performed with Student’s t-tests in SPSS 19.0. Results are expressed as means ± SD. P<0.05 indicates significance.

## Results

### Cells expressed fluorescence stably

Fluorescence was observed obviously in the cells in each well by in-vivo imaging system, and as the number of cells increased, the fluorescence intensity increased correspondingly (Fig 1A and B). It indicated that MDA-MB-231-luc cells could express luciferase stably.

**Fig 1.**
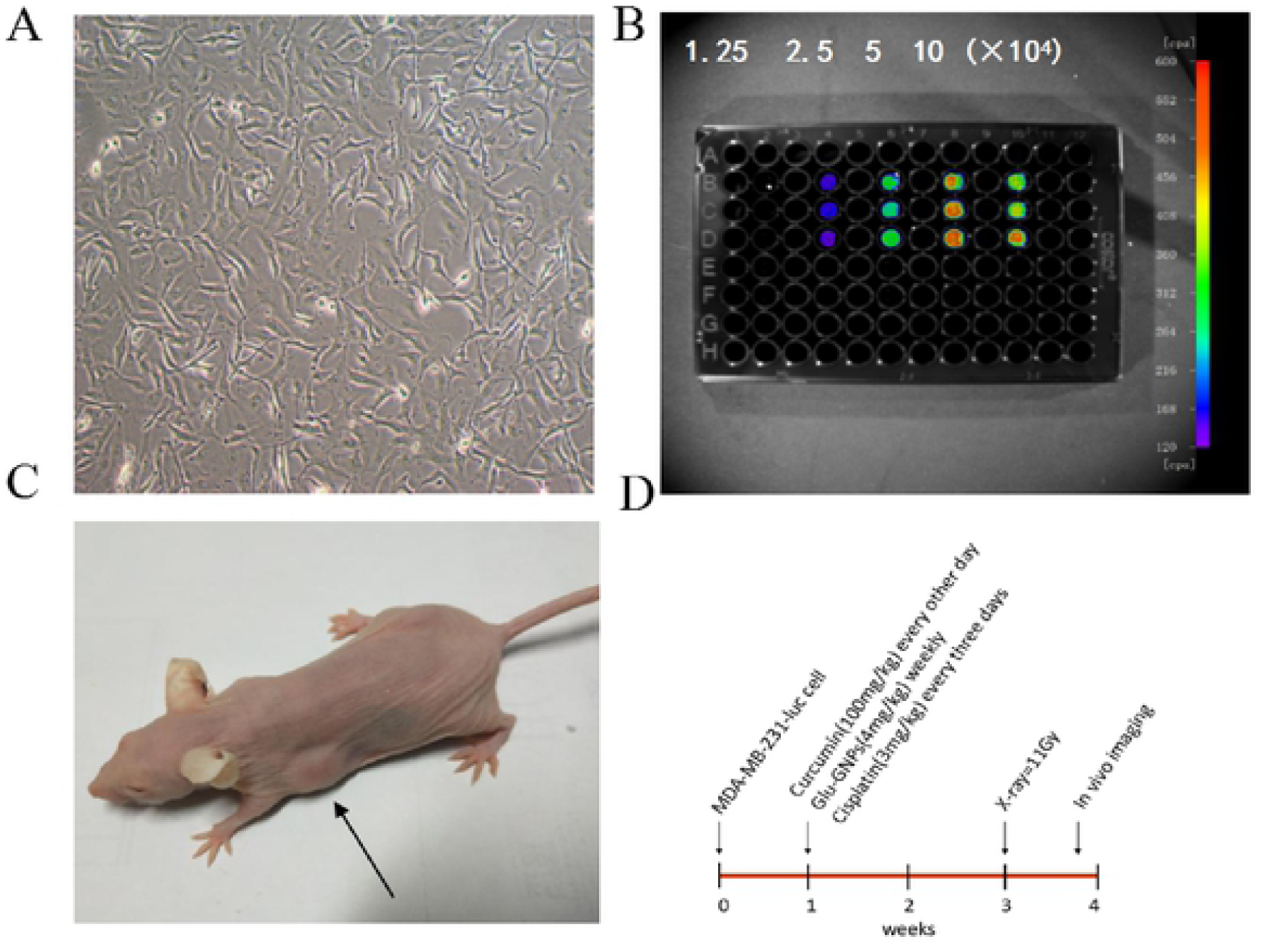
Cell culture and model establishment. (A) The image of MDA-MB-231-luc cell. (B) The fluorescence intensity of different number of cells(1.25×10^4^, 2.5×10^4^, 5× 10^4^ and 1× 10^5^ respectively)were detected in-vivo imaging system in the black opaque 96-well plate. (C) Breast tumor-bearing nude mice model with stable luciferase expression were successfully established 3-7 days after inoculation. (D) Schematic diagram of xenograft in nude mice. MDA-MB-231-luc cells were transplanted into the second pair of subcutaneous mammary glands on the left. When the volume of the tumor reached 100-200 mm^3^, curcumin in corn oil (2% DMSO), Glu-GNPs, and cisplatin in saline solution were administrated regularly. After 3 weeks of treatment, the tumor-bearing mice (IR, Glu-GNPs+IR, Cur+IR and Cur+Glu-GNPs+IR groups) were treated with irradiation (10Gy). Three days later, nude mice of each group were detected fluorescence intensity by in-vivo imaging system.

### Transplanted tumors in nude mice model was established successfully

Three to seven days after inoculated, nude mice formed tumors gradually and the tumor formation rate was 100% except for control group (Fig 1C). The shape of the tumor was close to sphere, and a few of them were irregular. Meanwhile, there was no significant change in body weight of tumor-bearing mice. When the volume of the tumor reached 100-200 mm^3^, the tumor-bearing mice were treated with different drugs for 3 weeks regularly (Fig 1D).

### Tumor growth was inhibited by curcumin

During treatment, the tumor volume was measured every 3 days and the body weight of tumor-bearing mice were measured every other week. Cisplatin was used as positive control. Change of tumor volume suggested that tumor volume in model group increased rapidly, while tumor growth in other treated groups was slower than that in model group, of which the cisplatin group grew slowly relatively and the tumor volume decreased less (p < 0.001). After irradiation treatment, the tumor volume of mice irradiated by X-ray decreased significantly, especially in Cur+Glu-GNPs+IR group (Fig 2A). Before radiotherapy treatment, the body weight of mice did not decrease obviously except for cisplatin group. After 3 weeks of medication, the tumor-bearing mice containing the irradiation group received X-ray treatment. After irradiation, the body weight of the mice decreased significantly and mice were generally in poor condition. Meanwhile, the body weight loss of mice in cisplatin group was still going on and two of them died, while there was no dead mice in curcumin group (Fig 2B). These results showed that curcumin can inhibit tumor growth without obvious toxic and side effects. The body weight of mice in Cur+Glu-GNPs+IR group decreased significantly, indicating that X-ray irradiation had a bad impact on the survival of mice. After sampled, the tumor tissue was weighed, and the tumor weight of each administration group was statistically different from that of the model group (Fig 2C and D). The tumor weight of cisplatin group, Cur+IR group and Cur+Glu-GNPs+IR group were reduced compared with model group. The results showed that curcumin combined Glu-GNPs along with radiation played a synergistic therapeutic effect on breast cancer tumor growth.

**Fig 2.**
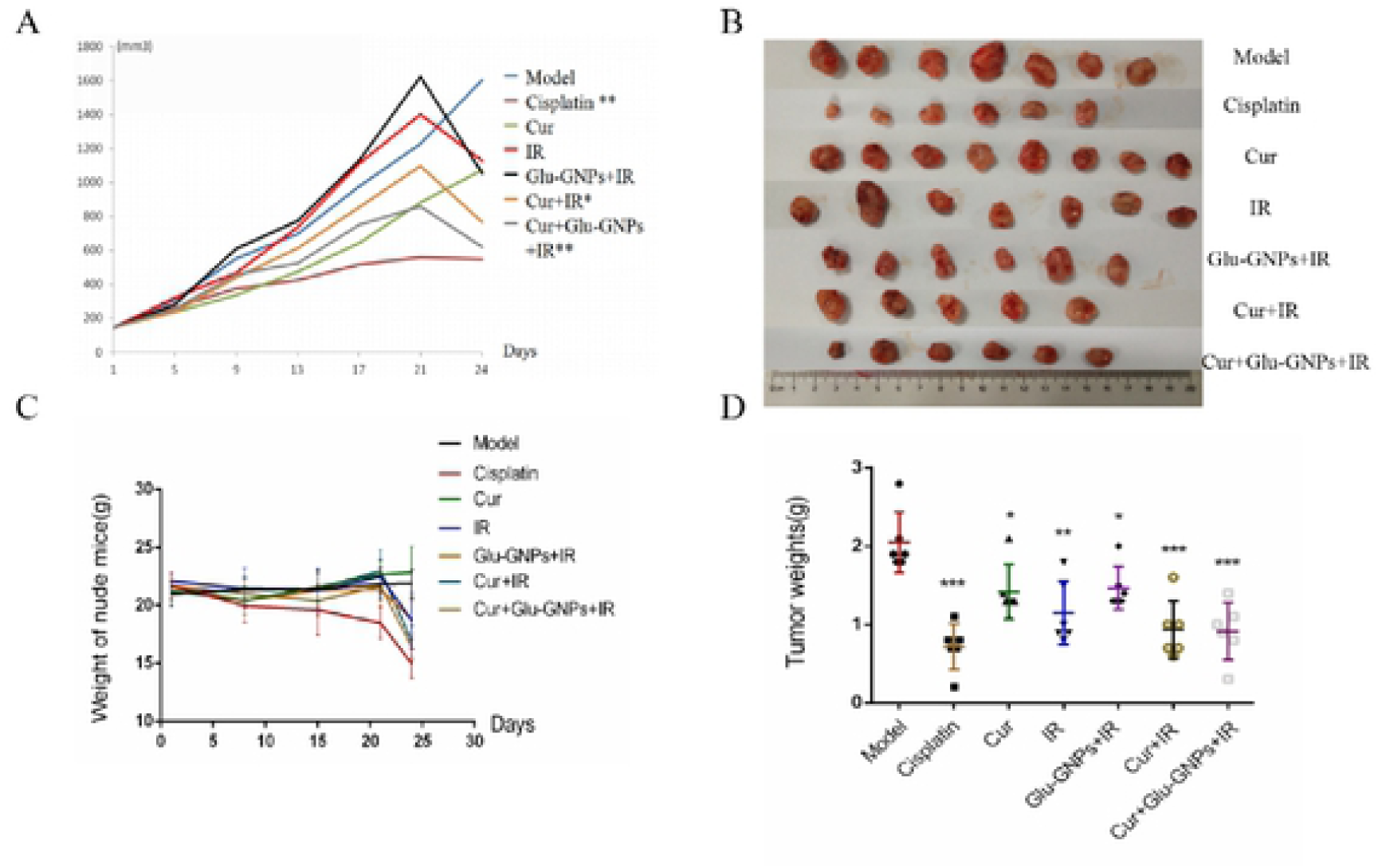
Tumor growth decreased after the treatment of curcumin, Glu-GNPs and radiation, alone and in combination. (A)Tumor volumes were measured every four days. The data was normalized to the model. P values were determined by the Student’s t test. ^*^P<0.05 and ^**^P<0.01 vs. Model group; (B) The weight of nude mice was measured weekly. (C) Images of transplanted tumor tissues. The tumor tissues were removed after treatment. (D)Tumor tissues weight was detected. The data was normalized to the Model group. Error bars indicate SD. ^*^P<0.05, ^**^P<0.01 and ^***^P<0.01 vs. Model group (Student’s t test).

### Changes of fluorescence intensity in-vivo imaging system

The fluorescence intensity of tumor growth was monitored 24 days after treatment, by in-vivo imaging system. We randomly selected two tumor-bearing mice in each group to detect their fluorescence intensity. The results showed that the fluorescence intensity of the model group was the highest among all tumor-bearing mice. Compared with the model group, the fluorescence intensity in cisplatin group and Cur+Glu-GNPs+IR group decreased significantly (Fig 3). It also showed that curcumin can inhibit tumor fluorescence intensity. And there was synergistic effect in curcumin combined Glu-GNPs along with radiation.

**Fig 3.**
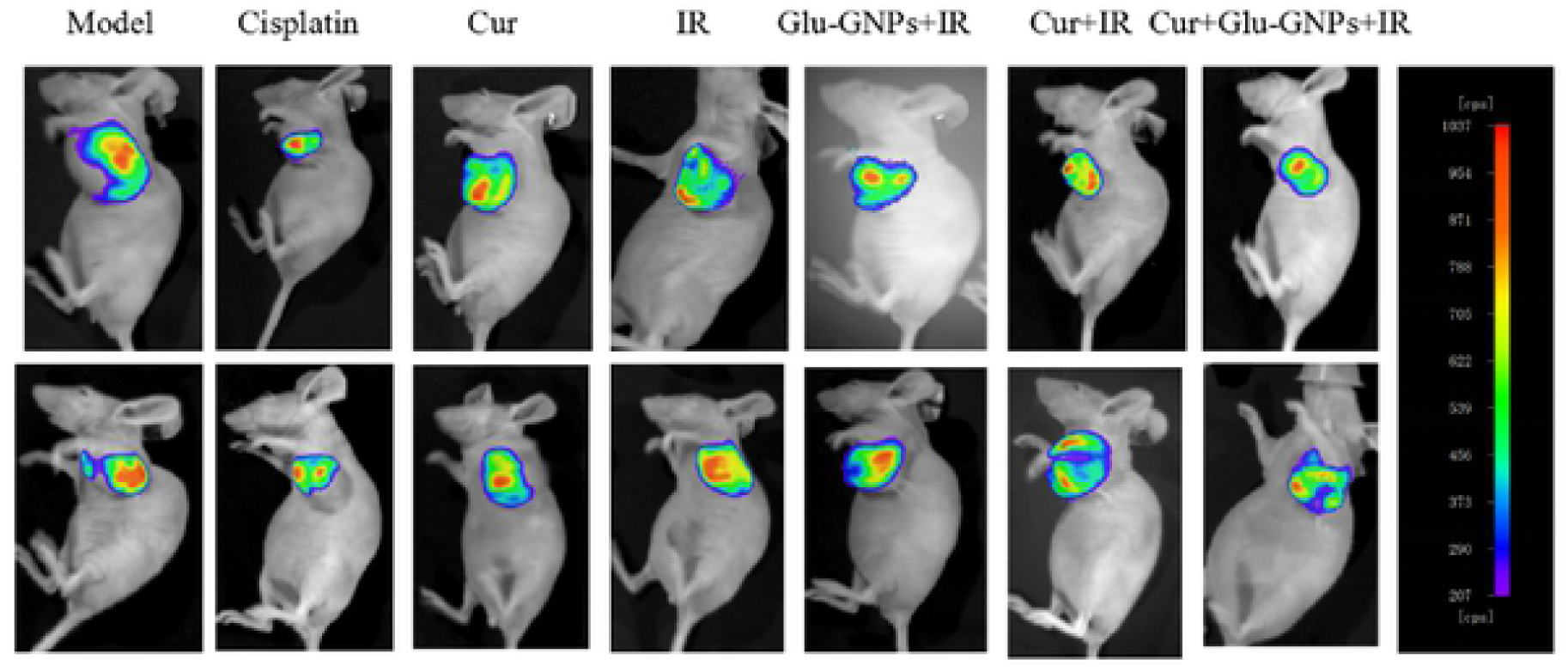
Changes of fluorescence intensity in-vivo imaging. After treatment, the fluorescence intensity of tumor-bearing mice injected luciferase substrate (150mg/kg) intraperitoneally in advance was monitored by in-vivo bioluminescence imaging system, and represented by the color. Mice were anesthetized by inhaling isoflurane. The fluorescence was observed under the same condition.

### Morphological changes of tumor tissues after treatment

After HE staining, the morphology of tumor tissue sections (4 μm) in each group were observed under microscope. The nucleus of tumor cells was large and heterogeneity, almost no cytoplasm, with more necrosis in the center of the tumor. The results showed that breast carcinoma xenograft was invasive ductal carcinoma with high malignancy. (Fig 4A).

**Fig 4.**
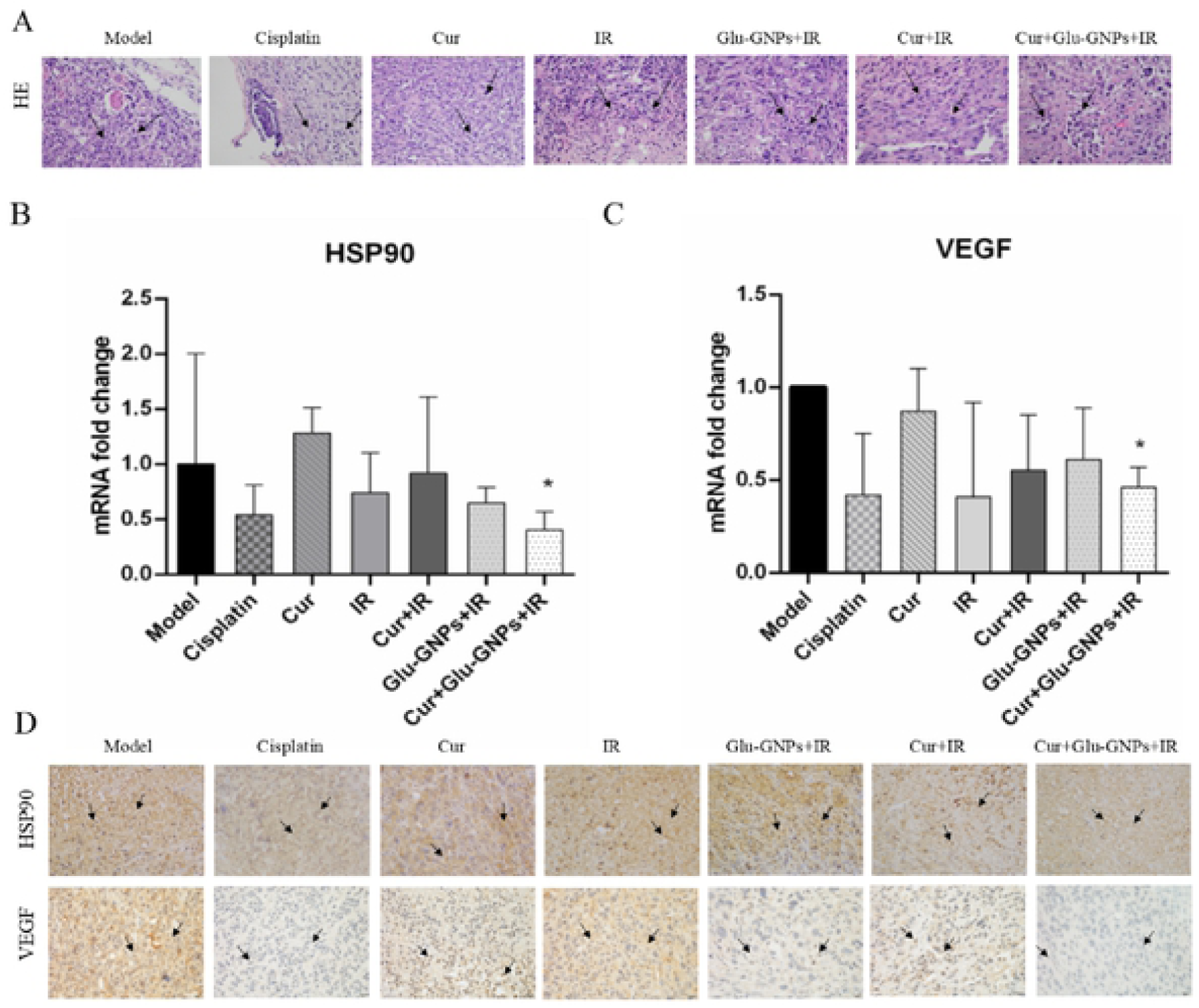
Detection of tumor tissue samples. (A)Representative images of HE staining(400×). HE staining depicts malignant degree of xenograft and form of tumor cells. The cells of tumor tissues had large and dark nuclei, almost no cytoplasm, with more necrosis in the center of the tumor. Arrows point to tumor cells. qRT-PCR analyses of the mRNA levels of *HSP90*(B) and *VEGF*(C) in tumor tissue. *β-actin* was used as reference gene. The data was analyzed with the mean ± SD of three times of independent experiments. ^*^ P<0.05 vs. Model group (Student’s t test). (D)Representative images of IHC staining(400×) of *HSP90* and *VEGF*. Tumor tissue was immunostained using DAB (brown) and Haematoxylin (blue) for nuclear counterstaining. Arrows point to nuclear expression of *HSP90* or *VEGF* in breast cancer cells respectively. Scale bars, 20 μm.

### mRNA level is regulated after treatment

The mRNA level of *VEGF* and *HSP90* was examined by qRT-PCR. Compared with model group, The mRNA level of VEGF [(Cisplatin (∼0.42-fold), Cur (∼0.87-fold), IR (∼0.87-fold), Cur+IR (∼0.55-fold), Glu-GNPs+IR (∼0.61-fold), and Cur+Glu-GNPs+IR(∼0.46-fold)] and HSP90[(Cisplatin(∼0.54-fold),Cur(∼1.28-fold),IR(∼0.74-fold),Cur+IR(∼0.92-fold),Gl u-GNPs+IR(∼0.65-fold),and Cur+Glu-GNPs+IR(∼0.40-fold)] were regulated(Fig. 6A and B).These results showed that curcumin and radiation can both down-regulate the expression level of *VEGF* in tumor tissue, especially Cur+Glu-GNPs+IR group significantly (P < 0.05). In addition, curcumin can down-regulate the mRNA level of *HSP90*. When curcumin combined Glu-GNP along with radiation, the expression level of *HSP90* decreased significantly compared with model group (P < 0.05) (Fig 4B and C). It was suggested that curcumin combined Glu-GNPs along with radiation may be related to the effective inhibition of *VEGF* and *HSP90* in breast tumor tissue.

### Immunohistochemical analysis of VEGF and HSP90 gene

Five areas were randomly selected under the microscope, of which brown was positive expression. The expression level of *VEGF* decreased in curcumin group, especially in cisplatin group and Cur+Glu-GNPs+IR group. *HSP90* expression in radiation group was higher than that in model group, but decreased after curcumin treatment (Fig 4D). Radiation can promote the expression of *HSP90*, while curcumin combined with irradiation can down regular the level of *HSP90*, protect normal cells and inhibit the growth of tumor cells. The results showed that curcumin can inhibit the growth of tumor vessels and had synergistic effect along with Glu-GNPs and X-ray therapy.

## Discussion

Curcumin is a plant polyphenol with effective activities and is also used as food in daily life. Nowadays, studies have reported that curcumin has anti-cancer activity, low toxicity and side effects. Our previous studies showed that curcumin has an effective inhibition on MDA-MB-231 and MCF-7 cells ^[22]^. It has been reported that curcumin combined with a variety of anti-cancer drugs can enhance the sensitivity of cancer cells, which has become a research direction. In addition, our previous studies had demonstrated that Glu-GNPs may be a promising radiosensitizer which exerted radiosensitizing effect on MDA-MB-231 adherent cells and stem cells. However, the influence of curcumin on radiosensitivity of breast carcinoma in vivo and the synergistic effect with Glu-GNPs are not clear.

In this study, we selected luciferase-labeled cell line MDA-MB-231-luc, and established transplanted tumor model by subcutaneous inoculated of cells into the underarm of mice, with a tumor formation rate of 100%. Then nude mice were treated for 3 weeks, cisplatin was used as positive drug. According to the growth rate of tumor volume and the body weight of mice, the curative effect of drugs was preliminarily evaluated. We found that cisplatin was effective in the treatment process, but the body weight loss of mice in cisplatin group was obvious. Curcumin-containing group could not only inhibit tumor growth, but also had little effect on the body weight of tumor-bearing mice.

After that, we observed the tumor fluorescence intensity of each group of tumor-bearing mice through in-vivo imaging assay. The results showed that the tumor fluorescence intensity in the model group was the highest, while in the Cur(100mg/kg) group, Cur+IR group and Cur+Glu-GNPs group, the fluorescence intensity is relatively weakened. The decrease was the most obvious in Cur+Glu-GNPs+IR group (Tumor growth rate=5.92%) and cisplatin group (Tumor growth rate=5.31%).Therefore, it can be concluded that curcumin can effectively inhibit the growth rate of transplanted tumor, and combined Glu-GNPs along with X-ray has synergistic therapeutic effect.

Studies have shown that its radiotherapy resistance mechanism is that blood vessels are irregularly distributed in tumor tissue, blood circulation is blocked, hypoxic areas exist in tumor tissue, hypoxic cells have high radiation tolerance and DNA damage of tumor stem cells ^[23]^. Therefore, the combination of drugs and irradiation may be sensitive to tumor cells. In this study, we found that curcumin combined with Glu-GNPs can significantly reduce *VEGF* mRNA and protein levels (p < 0.05), and it can reduce HSP90 production in tumor tissue. *VEGF* is an angiogenesis factor in mice and plays an important role in tumor growth. The level of *VEGF* can be used as a marker to assess tumor status. *HSP90* is a kind of stress protein, which can rise sharply and promote the growth of tumors when stimulated by the environment. The results suggested that curcumin can inhibit the generation of blood vessels and *HSP90* in breast cancer transplanted tumor, thus inhibiting the growth of tumor and promoting the killing effect of radiation on tumor cells.

## Acknowledgments

The author would like to thank the Animal Ethics Committee of Guangzhou University of Chinese Medicine for its support and the experimental platform provided by School of Basic Medical Science.

## Author Contributions

Data curation: Mengjie Li, Tingting Guo, Chenxia Hu.

Formal analysis: Mengjie Li, Tingting Guo, Yujian Wu.

Investigation: Mengjie Li, Tingting Guo, Ke Yang, Chenxia Hu.

Methodology: Mengjie Li, Tingting Guo, Chenxia Hu.

Project administration: Chenxia Hu

Software: Yujian Wu, Ke Yang, Jiayi Lin, Mengjie Li.

Supervision: Chenxia Hu.

Writing-original draft: Mengjie Li, Yujian Wu.

Writing-review & editing : Mengjie Li, Jiayi Lin, Tingting Guo, Ke Yang, Chenxia Hu.

